# CRISPRi is not strand-specific and redefines the transcriptional landscape

**DOI:** 10.1101/156950

**Authors:** Françoise S. Howe, Andrew Russell, Afaf El-Sagheer, Anitha Nair, Tom Brown, Jane Mellor

## Abstract

CRISPRi, an adapted CRISPR-Cas9 system, is proposed to act as a strand-specific roadblock to repress transcription in eukaryotic cells using guide RNAs (sgRNAs) to target catalytically inactive Cas9 (dCas9) and offers an alternative to genetic interventions for studying pervasive antisense transcription. Here we successfully use click chemistry to construct DNA templates for sgRNA expression and show, rather than acting simply as a roadblock, binding of sgRNA/dCas9 creates an environment that is permissive for transcription initiation and termination, thus generating novel sense and antisense transcripts. At *HMS2* in *Saccharomyces cerevisiae*, sgRNA/dCas9 targeting to the non-template strand results in antisense transcription termination, premature termination of a proportion of sense transcripts and initiation of a novel antisense transcript downstream of the sgRNA/dCas9 binding site. This redefinition of the transcriptional landscape by CRISPRi demonstrates that it is not strand-specific and highlights the controls and locus understanding required to properly interpret results from CRISPRi interventions.

## INTRODUCTION

Eukaryotic genomes are pervasively transcribed but the effect of this inter– and intragenic transcription is not fully understood (Mellor, Woloszczuk, & Howe, 2016). Of particular interest is the function of antisense transcription, which alters the chromatin in the vicinity of sense promoters (Lavender et al., 2016; Murray et al., 2015; Pelechano & Steinmetz, 2013), and is associated with repression, activation or no change in levels of the corresponding sense transcript (Murray et al., 2015). Discerning the mechanism by which antisense transcription functions in gene regulation is confounded by the difficulty in ablating antisense transcription without direct genetic intervention (Bassett et al., 2014). A new approach, CRISPR interference (CRISPRi) (Qi et al. 2013), can circumvent these issues by avoiding the need to manipulate endogenous DNA sequences. An endonucleolytically dead version of Cas9 (dCas9) is recruited by single base pairing guide RNAs (sgRNAs) targeting the non-template (NT) DNA strand, where it blocks transcription strand-specifically at the loci tested (Lenstra, Coulon, Chow, & Larson, 2015; Qi et al., 2013). In a further adaptation, dCas9 fusion with transcriptional repressors or activators allows both negative and positive regulation of transcription respectively (Gilbert et al., 2013, 2014), but the strand-specificity is lost and thus is not suitable for strand-specific repression of antisense transcription. The sgRNA compenent of the CRISPRi system consists of two regions: a constant region (82 nt) that binds to dCas9 and a variable region (20 nt) that is responsible for targeting. The modular nature of the sgRNA lends itself to synthetic copper(I)-catalysed alkyne-azide cycloaddition (CuAAC) click chemistry (El-Sagheer & Brown, 2010). The stable artificial triazole DNA linker generated is biocompatibile, being read and accurately copied by DNA and RNA polymerases (Birts et al., 2014; El-Sagheer & Brown, 2011; El-Sagheer, Sanzone, Gao, Tavassoli, & Brown, 2011). We show that click chemistry is an efficient method for sgRNA construction and that when combined with dCas9, these sgRNAs are as effective as sgRNAs produced from chemically synthesised full-length DNA templates at reducing levels of transcripts *in vivo.* However, it is still not completely understood how the CRISPRi system functions strand-specifically or why sgRNAs must target the non-template strand.

Here we use the CRISPRi system to study the effect of blocking antisense transcription at loci with well characterised sense:antisense transcript pairs. We have previously used a promoter deletion of the antisense transcript *SUT650* at the *HMS2* locus to show that *SUT650* represses *HMS2* sense transcription (Nguyen et al., 2014). Now we use CRISPRi to examine the effects of blocking *SUT650* antisense transcription to ask (i) whether *SUT650* represses *HMS2* sense transcription *without* a genetic intervention and (ii) whether CRISPRi is strand-specific using, in addition to *HMS2,* an engineered *GAL1* gene with a well characterised antisense transcript (Murray et al., 2012, 2015).

The main conclusion from this study is that CRISPRi at the *HMS2* locus is not fully strand-specific and results in (i) premature termination of the sense transcript and (ii) initiation of a new unstable antisense transcript in the vicinity of the sgRNA binding site. As transcription from this new antisense initiation site extends in to the *HMS2* promoter, there is no net change in *HMS2* sense transcript levels. Thus CRISPRi redefines the transcriptional landscape at *HMS2.* This suggests that routine use of CRISPRi for gene expression analysis will require rigorous analysis of transcript integrity and function before conclusions can be drawn.

## RESULTS & DISCUSSION

### DNA templates for sgRNA production can be made using click chemistry

CRISPRi-mediated transcriptional repression requires co-expression of a mature sgRNA and dCas9 (Fig 1A). A small library of single-stranded DNAs, comprised of templates for sgRNA variable regions, were joined to the constant region using click chemistry (Fig. 1B) and used successfully as templates for PCR, with no significantly different efficiencies when compared to control full-length synthesised oligonucleotides (Fig. 1C). The PCR products were inserted in place of a *URA3* selection cassette in the endogenous *snR52* locus for expression of a transcript that is then processed to form the mature nuclear-retained sgRNA (Fig. 1A). Levels of dCas9 protein were uniform between strains (Fig. 1 - Figure Supplement 1). Neither insertion of *URA3* into snR52 nor dCas9 expression in the control strains affected growth rate, although strains expressing some sgRNAs grew more slowly indicating a physiological effect (Fig. 1D).

**Figure 1.**
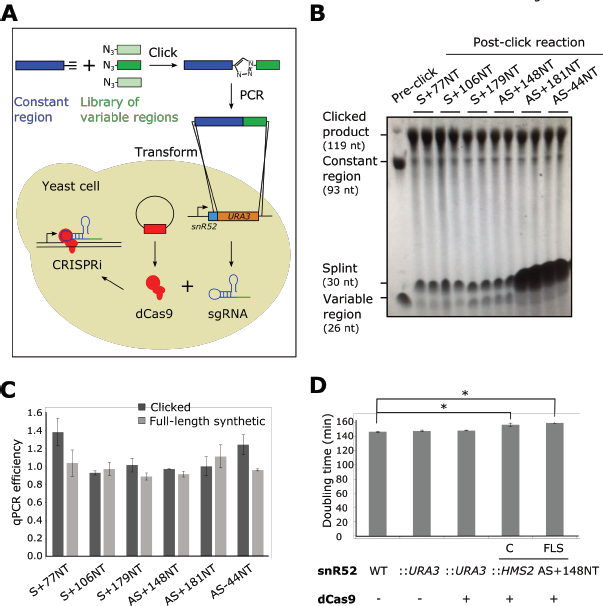
Experimental strategy for sgRNA template synthesis and incorporation into yeast. **A.** An outline of the experiment. Templates for sgRNA production were generated using a click chemistry reaction between a single-stranded DNA oligonucleotide with a 3’ alkyne group encoding the constant region (dark blue) and a number of single-stranded DNA oligonucleotides with 5’ azido groups encoding the different variable regions (green). The resulting single-stranded DNA oligonucleotide was purified, amplified by PCR and transformed into yeast to replace the *URA3* selection cassette that had previously been introduced into the endogenous *snR52* locus. Correct insertion was confirmed by Sanger sequencing. The expression of the sgRNA is driven from the endogenous RNA polymerase III *snR52* promoter and processed to produce a mature sgRNA. The sgRNA couples with enzymatically dead Cas9 (dCas9, red) expressed under the control of the *TDH3* promoter off a plasmid to block transcription. See Methods for more detail. **B.** A polyacrylamide gel visualised by UV shadowing reveals a high efficiency of the click reaction for all the *HMS2* constructs (see Figures 2E and Figure 3 - Figure Supplement 1 for positions of the sgRNA binding sites). The sizes of the variable region, DNA splint, constant region and clicked product are indicated. **C.** PCR efficiencies of the clicked and full-length synthetic DNA oligonucleotides as measured by qPCR with a serial dilution series. N=3, errors are standard error of the mean (SEM), all differences are not significant p>0.05. **D.** Doubling times (min) of the indicated yeast strains grown in complete synthetic media minus leucine. C, strains constructed using clicked oligonucleotides; FLS, strains constructed using full-length synthetic oligonucleotides. N=10, errors are SEM, * p<0.05.

### CRISPRi represses the production of antisense transcripts at *HMS2* and *GAL1*

CRISPRi represses transcription when sgRNAs/dCas9 are targeted to the non-template (NT) strand next to a protospacer adjacent motif (PAM) (Qi et al., 2013) (Fig. 2A). Firstly, we used an engineered version of *GAL1* that has a stable antisense transcript *(GAL1* AS) initiating within an *ADH1* terminator inserted into the *GAL1* coding region (Murray et al., 2012) (Fig. 2B). *GAL1* AS is present in cells grown in glucose-containing media when the *GAL1* gene is repressed and is reduced as cells are switched into galactose-containing media and *GAL1* sense is induced (Fig. 2C lanes 1-3). We designed sgRNAs adjacent to two PAM sequences on the non-template strand near the antisense transcription start site (TSS) (AS+28NT and AS+112NT) and a third strand-specificity control sgRNA on the template strand in this region (AS+93T) (Fig. 2B). Only sgRNA AS+112NT/dCas9 caused significant (p=0.016) reduction in *GAL1* antisense transcript levels, as assessed by Northern blotting (Fig. 2C,D).

**Figure 2.**
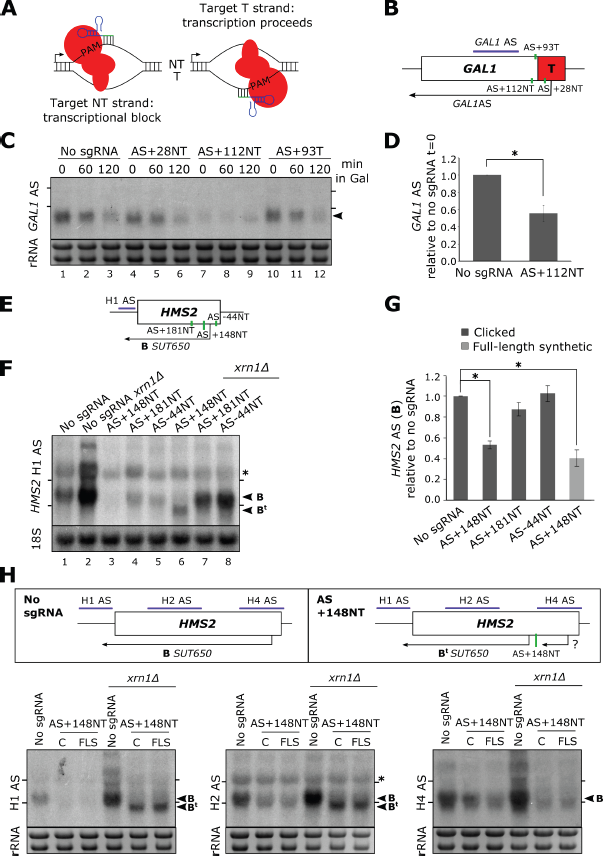
CRISPRi reduces antisense transcripts at *GAL1* and *HMS2*. **A.** A schematic demonstrating the previously reported requirement of non-template strand targeting by the sgRNA/dCas9 complex for strand-specific transcriptional repression. Arrows indicate the direction of transcription. NT, non-template; T, template; red, dCas9; PAM, protospacer adjacent motif; green/blue, sgRNA. **B.** A schematic of the engineered *GAL1* locus showing the site of insertion of the *ADH1* terminator (T, red box), the transcription start site for the stable *GAL1* antisense transcript and the positions of the designed sgRNAs (green vertical lines) targeting the non-template (AS+28NT, AS+112NT) and template (AS+93NT) strands with respect to antisense transcription. The position of the Northern blotting probe detecting the *GAL1* AS transcript is shown in purple. **C.** A Northern blot showing the reduction in *GAL1* antisense transcript (black arrowhead) in the strain expressing sgRNA AS+112NT/dCas9 relative to the no sgRNA control *(GAL1:ADH1t snR52::URA3* with dCas9). sgRNAs AS+28NT and AS+93T do not alter *GAL1* AS levels. Samples were taken from cells grown in glucose (t=0) and at the indicated times after transfer to galactose-containing media (min). Positions of the rRNA are indicated by the short horizontal lines. Ethidium bromide-stained rRNA is used as loading control. **D.** Quantification of Northern blotting for the *GAL1* AS transcript at t=0 in the control no sgRNA strain and the strain with sgRNA AS+112NT. N=6, errors are SEM, * p<0.05. **E.** A map of the *HMS2* locus showing the *HMS2* AS transcript, *SUT650* (black arrow, transcript B) and positions of the three sgRNAs targeting *SUT650* (green vertical lines). The position of the Northern blotting probe to detect *SUT650* (H1 AS) is shown by the purple line. **F.** A Northern blot probed with *HMS2* antisense probe H1 (see schematic in (E)) showing the level of *SUT650* (black arrowhead, transcript B) in the no sgRNA control *(snR52::URA3* with dCas9) strain and strains expressing the indicated antisense-targeting sgRNAs. Deletion of *XRN1* allows detection of a shorter antisense transcript (B^t^) in the strain expressing AS+148NT. Positions of the rRNA are indicated by the short horizontal lines. *represents cross-hybridisation with the 26S rRNA. A blot probed for the 18S rRNA is used as loading control. **G.** Quantification of the level of *SUT650* (transcript B) reduction in the strains expressing each of the 3 antisense sgRNAs relative to the control no sgRNA strain. N=4-8, errors represent SEM, *p<0.05. Click-linked and full-length synthetic sgRNA AS+148NT templates behave similarly. **H.** Northern blots with a series of *HMS2* antisense-specific probes. A new shorter antisense transcript (B^t^) can be detected upon *XRN1* deletion in the strains expressing sgRNA AS+148NT/dCas9. Two clones of the CRISPRi strains produced using clicked (C) or full-length synthetic (FLS) oligos for strain construction are shown. Positions of the antisense-specific probes (purple) and the site of sgRNA AS+148NT/dCas9 binding (green vertical line) are shown in the schematic. Positions of the rRNA on the Northern blot are indicated by the short horizontal lines. *represents cross-hybridisation with the 26S rRNA. Ethidium bromide-stained rRNA is used as loading control.

Next, we examined the *HMS2* locus which has a stable antisense transcript, *SUT650,* initiating within the 3’ coding region of *HMS2* (Nguyen et al., 2014; Pelechano, Wei, & Steinmetz, 2013). We designed three SUT650-targeting sgRNAs, one acting as a control located 44 nucleotides upstream of the *SUT650* TSS on the non-template strand (AS-44NT), and two sgRNAs, AS+148NT and AS+181NT, targeting sequences downstream of the TSS (Fig. 2E). AS+148NT/dCas9 significantly (p=0.011) reduced *SUT650* (Fig. 2F lanes 1&3, 2G) and this repression was comparable for the clicked and control full-length synthetic constructs (Fig. 2G). However, there was no obvious effect of the other two sgRNAs/dCas9 on *SUT650* levels (Fig. 2F lanes 1, 4&5, 2G). CRISPR efficiency has been linked to nucleosome occupancy (Horlbeck, Witkowsky, et al., 2016; Isaac et al., 2016) but we found no correlation between the level of repression and the nucleosome occupancy (Knight et al., 2014) at each of the *SUT650* sgRNA target regions (Fig. 2 - Figure Supplement 1). However, other factors such as sequence determinants can also influence repression (Horlbeck, Gilbert, & Villalta, 2016). These results at *GAL1* and *HMS2* confirm CRISPRi is suitable for reducing levels of antisense transcripts in yeast but controls are needed for each sgRNA designed to ensure that repression has been achieved.

### CRISPRi repression of *SUT650* induces a new shorter antisense transcript at *HMS2*

At *HMS2,* sgRNA AS+148NT/dCas9 was able to reduce *SUT650* levels significantly (Fig. 2F,G). Since *SUT650* is a substrate for the major cytoplasmic 5’-3’ exonuclease Xrn1 (Fig. 2F, lanes 1&2), we investigated whether the CRISPRi-induced *SUT650* reduction was due to direct repression of antisense transcription and/or a reduction in *SUT650* stability. Whilst *SUT650* was not stabilised upon *XRN1* deletion in the strain expressing AS+148NT/dCas9, supporting previous studies that CRISPRi operates via a transcriptional block (Qi et al., 2013) (Fig. 2F, lanes 3&6, transcript B), we detected a shorter antisense transcript (Fig 2F, lane 6, transcript B^t^). To map transcript B^t^, we used a series of Northern blotting probes across the locus (Fig. 2H). With antisense-specific probes H1 and H2 in the *HMS2* gene promoter and 5’ coding region respectively, we could detect transcript B^t^, indicating that it extends into the *HMS2* sense promoter. However, probe H4 at the 3’ end of the *HMS2* gene was unable to hybridise to B^t^, indicating that B^t^ initiates upstream of this site. So whilst CRISPRi successfully blocked the original *SUT650,* a new Xrn1-sensitive unstable antisense transcript (B^t^) was created. This is potentially due to changes in the chromatin structure caused by AS+148NT/dCas9 binding to this region (Horlbeck, Witkowsky, et al., 2016; Isaac et al., 2016), creating an environment that is permissive for transcription initiation (Murray et al., 2012). We note that using probe H4 we were also unable to detect *SUT650* initiating from its WT start site but terminating at the AS+148NT binding site, due to its small size (~148 nt).

### CRISPRi-induced *GAL1* AS repression does not affect the *GAL1* sense transcript

We tested whether *GAL1* AS repression by AS+112NT/dCas9 affected the *GAL1* sense transcript. Previously we mutated a TATA-like sequence to ablate antisense transcription but observed no difference in *GAL1* sense transcript levels in the population (Murray et al., 2015). Using CRISPRi, we also observed no significant change in *GAL1* sense transcript levels or size (Fig. 3A,B). To support a strand-specific transcriptional block of AS transcription, *GAL1* sense transcript polyA site usage was unaffected by sgRNA AS+112NT/dCas9 (Fig. 3C). Furthermore, *XRN1* deletion in this strain also did not affect polyA site usage, ruling out a partial double-stranded transcriptional block and subsequent destabilisation of the resulting truncated sense transcript (Fig. 3C).

**Figure 3.**
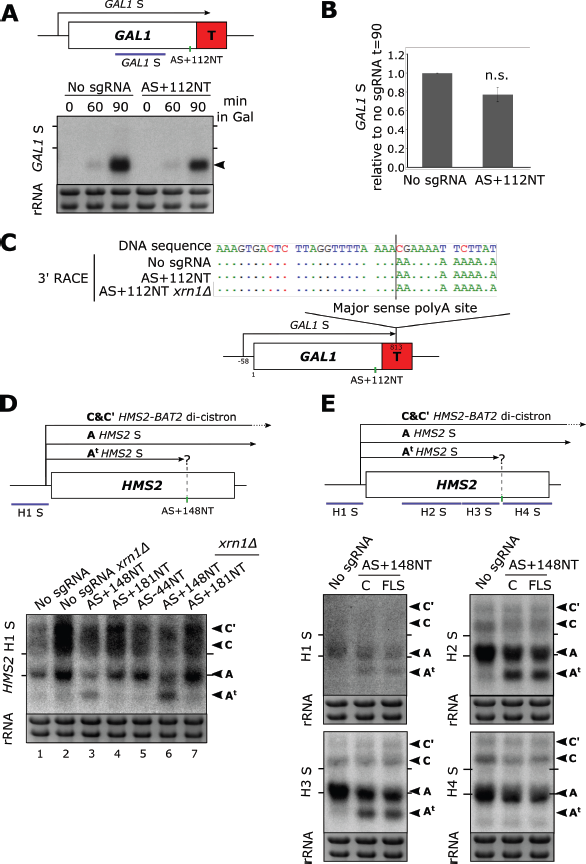
HMS2 and GAL1 sense transcripts are differently affected by blocking of their respective antisense transcripts. **A.** A Northern blot probed for the *GAL1* sense transcript (black arrow) in the control strain with no sgRNA *(GAL1 ADHlt snR52::URA3* with dCas9) and in the strain expressing sgRNA AS+112NT/dCas9. Samples were taken from cells grown in glucose (t=0) and at the indicated times after transfer to galactose-containing media (min). Ethidium bromide-stained rRNA is used as a loading control. **B.** Quantification of the Northern blot shown in (A) at t=90. N=4, error bars represent SEM, change is statistically non-significant (n.s.) p=0.062. **C.** A schematic showing the results of *GAL1* sense transcript 3’ end mapping by RACE in the strains indicated. All distances are shown relative to the sense TSS. Transcript cleavage and polyadenylation occurs beyond the position of the *GAL1* AS– blocking AS+112NT sgRNA shown (green vertical line). **D.** A Northern blot probed with *HMS2* sense probe H1 (purple) showing the level of *HMS2* sense (black arrowhead, transcript A) in the no sgRNA control *(snR52::URA3* with dCas9) strain and strains expressing the indicated antisense-targeting sgRNAs. A truncated sense transcript (A^t^) was also detected in the strain expressing AS+148NT. Positions of the rRNA are indicated by the short horizontal lines. Ethidium bromide-stained rRNA is used as loading control. **E.** Northern blots with a series of *HMS2* sense-specific probes detecting the regions indicated in the schematic. The position of the sgRNA AS+148NT/dCas9 binding site is shown (green vertical line). Positions of the rRNA are indicated by the short horizontal lines. Ethidium bromide-stained rRNA is used as loading control.

### CRISPRi-induced *SUT650* repression truncates the *HMS2* sense transcript

Previous work shows that reducing *SUT650* transcription increases *HMS2* sense transcript levels (Nguyen et al., 2014). However, blocking *SUT650* by AS+148NT/dCas9 did not similarly increase *HMS2* sense levels (Fig. 3D, lanes 1&3). To our surprise, in addition to the full-length *HMS2* sense transcript (A), we detected a considerably shortened *HMS2* sense transcript (A^t^). Since the combined levels of the truncated and full-length transcripts are similar to those in the strain not expressing an sgRNA, independent of the presence of *XRN1* (Fig. 3D, lanes 1&3 or 2&6), we hypothesised that the AS+148NT/dCas9 complex bound within the *HMS2* coding region may be causing partial premature sense transcription termination, leading to transcript A^t^. Thus, we mapped the 3’ end of *HMS2* S by Northern blotting using a series of strand-specific probes across the locus (Fig. 3E). Both transcripts A and A^t^ could clearly be detected using the probes upstream of the AS+148NT/dCas9 binding site (probes H1, H2, H3) but the truncated transcript was undetectable using probe H4 downstream of this site. Thus the transcriptional block caused by AS+148NT/dCas9 is not strand-specific and redefines the transcriptional landscape at the *HMS2* locus by creating a chromatin environment that is suitable for both transcription initiation and termination. As an additional control, we designed three sgRNAs to block the *HMS2* sense transcripts (S+77NT, S+106NT, S+179NT) and monitored their effects on both *HMS2* sense and antisense transcripts (Fig. 3 - Figure Supplement 1). However, we observed no effect on either levels or integrity.

### CRISPRi-induced antisense repression at *HMS2* and *GAL1* is not as efficient as previous methods

We compared the repression efficiency of CRISPRi with our previously used genetic interventions. At *GAL1,* mutation of a TATA-like sequence within the *ADH1* terminator greatly reduced the level of the *GAL1* antisense transcript (Fig. 4A,B) (Murray et al 2015). *XRN1* deletion in this strain only slightly increased *GAL1* AS levels, suggesting that most repression was at the level of transcription rather than altering transcript stability (Fig. 4B,C). By contrast, *XRN1* deletion in the strain expressing AS+112NT/dCas9 did result in some upregulation of *GAL1* AS (Fig. 4B,C), indicating that antisense transcript repression was not as great in this strain. Unlike at *HMS2* (Fig. 2F,G), this stabilised *GAL1* AS transcript was the same size as in the control and so likely represents stabilisation of the transcripts escaping repression rather than novel transcripts.

**Figure 4.**
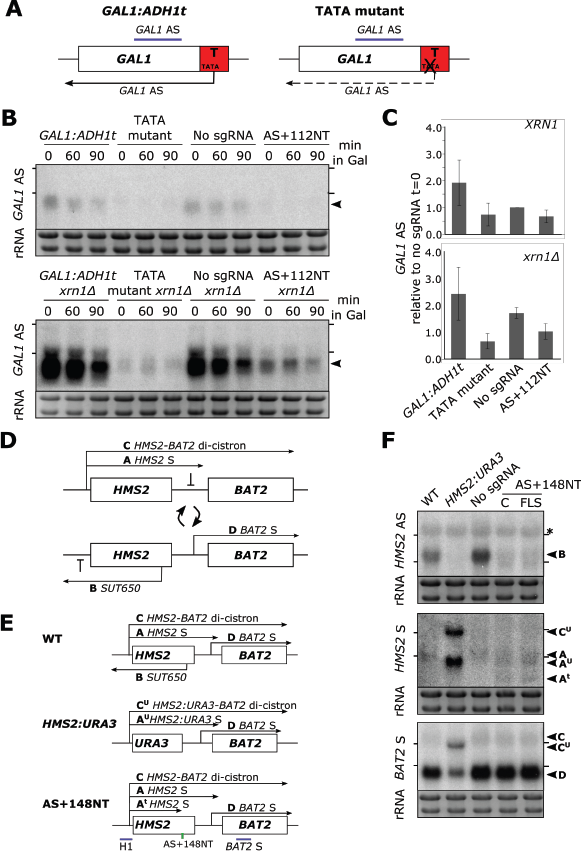
CRISPRi block of *SUT650* and *GAL1* AS is not as efficient as previous genetic interventions. **A.** A schematic showing the previously used genetic intervention to repress the *GAL1* AS transcript (Murray et al., 2015). In the TATA mutant construct, a TATA box within the engineered *ADH1* terminator (T) is scrambled and the level of *GAL1* antisense transcript is reduced. **B.** A Northern blot probed for the *GAL1* antisense transcript (black arrowhead) comparing the two methods of reducing antisense at this locus in the presence (top blot) or absence (bottom blot) of *XRN1.* All samples were run on the same gel and the blots were exposed to film for the same time. Samples were taken from cells grown in glucose (t=0) and at the indicated times after transfer to galactose-containing media (min). Positions of the rRNA are indicated by the short horizontal lines. Ethidium bromide-stained rRNA is used as loading control. **C.** Quantification of the Northern blot in (B) at t=0. N=2, errors represent the SEM. **D.** A schematic of the *HMS2-BAT2* locus, showing the *HMS2* sense and antisense transcripts (A and B), the *HMS2-BAT2* di-cistronic transcripts (C and C’) and the *BAT2* sense transcript (D). Cells switch between sense– and antisense-dominant states in which the indicated transcripts are produced (Nguyen et al., 2014). **E.** The *HMS2-BAT2* locus and associated transcripts in the WT strain and strains that repress the *SUT650* transcript. In the *HMS2:URA3* strain, the entire coding region of *HMS2* is replaced by that of *URA3.* **F.** Northern blots comparing the efficiency of *SUT650* reduction by replacement of the *HMS2* coding region with that of *URA3* versus CRISPRi and the relative effects of these two strategies on the *HMS2* sense and *BAT2* transcripts shown in (E). The WT (BY4741) and *HMS2.URA3* strains have been transformed with plasmid pRS315 to allow for growth on complete synthetic media lacking leucine and thus comparisons with the CRISPRi strains. Positions of the rRNA are indicated by the short horizontal lines. *represents cross-hybridisation with the 26S rRNA. Ethidium bromide-stained rRNA is used as loading control.

Next we compared the efficacy of AS+148NT/dCas9 with previous experiments to ablate *SUT650*, where we replaced the entire *HMS2* coding region, including the antisense TSS with the *URA3* coding region *(HMS2:URA3)* (Nguyen et al., 2014). This successful block of *SUT650* transcription increased levels of the *HMS2* sense transcript, the di-cistronic *HMS2-BAT2* transcript and consequently decreased levels of the downstream gene, *BAT2,* due to transcriptional interference (Fig. 4D-F). Whilst CRISPRi resulted in a similar decrease in *SUT650* as *HMS2:URA3,* neither the *HMS2, BAT2* nor *HMS2-BAT2* di-cistronic transcripts were altered (Fig. 4F). The discovery that whilst AS+148NT/dCas9 represses *SUT650,* a new unstable *HMS2* antisense transcript is induced and the sense transcript is prematurely terminated could explain why blocking *SUT650* using CRISPRi and the *URA3* gene body replacement strategies did not give the same results (Fig. 4F). Thus CRISPRi is not as effective as a genetic mutation in reducing levels of either the *GAL1* or *HMS2* AS transcripts.

## Concluding remarks

Although CRISPRi has been used to strand-specifically repress antisense transcription at *GAL10*(Lenstra et al., 2015) and *GAL1* (this work), here we demonstrate that this is not true at *HMS2.* This may reflect the vastly different transcription levels for the galactose inducible genes (high) compared to *HMS2* (low). Furthermore, a study in human K562 cells found that CRISPRi-induced repression did not correlate with which strand was targeted at 49 genes, although the mechanism behind this observation was not investigated (Gilbert et al., 2014).

This work extends our previous hypothesis (Nguyen et al., 2014), allowing us to propose that, rather than acting as a roadblock, the binding of the sgRNA/dCas9 complex at *HMS2* creates a new chromatin environment, which is permissible for transcription initiation and termination. Thus, new transcription units are generated that result in novel sense and antisense transcripts of varying stabilities that are therefore not always detectable. This redefinition of the transcriptional landscape highlights the levels of controls and locus understanding needed before results from using CRISPRi can be interpreted.

## AUTHOR CONTRIBUTIONS

Conception and design: FSH, AE-S, TB, JM; Acquisition of data: FSH, AR, AN; Analysis and interpretation of the data: FSH, AR, AN, JM; Drafting or revising the manuscript: FSH, TB, JM; Contributed unpublished, essential data, or reagents: AE-S.

## ACKNOWLEDGEMENTS

The authors thank Jack Feltham for assistance with sgRNA template construction.

## MATERIALS AND METHODS

### sgRNA design

PAM sequences within 200 bp of the TSS in question were identified and the 20 bp immediately adjacent to these were used to design the variable regions of the sgRNAs. These sequences were run through an off-spotter algorithm (https://cm.jefferson.edu/Off-Spotter//) to minimise any off-target effects (Pliatsika & Rigoutsos, 2015). The sequence for the engineered constant region and *SUP4* terminator were taken from (Dicarlo et al., 2013; Mali et al., 2013). The *HMS2* sgRNA templates were synthesised as two DNA oligonucleotides to be joined by click chemistry, with the majority of the constant region on one oligonucleotide that could be used for all reactions. To increase the efficiency of the click reaction, the click linkage was designed between a dCpT dinucleotide within the Cas9 handle region. The sequences for the *HMS2* and *GAL1* sgRNAs are shown in Supplementary File 1. Full-length sgRNA templates were also synthesised for the *GAL1* sgRNAs and as a control for the *HMS2* click chemistry constructs (Eurofins Genomics).

### Click chemistry

Single-stranded DNA oligonucleotides complementary to the constant and variable sgRNA regions were synthesised by standard phosphoramidite oligonucleotide synthesis with 3’-alkyne or 5’-azido modifications respectively (El-Sagheer et al., 2011) (Supplementary File 1). The click chemistry reaction was performed with a Cu^1^ catalyst and 24-30 nt splint DNAs (El-Sagheer et al., 2011). PCR was performed for second strand synthesis, amplification and incorporation of sequences homologous to the site of insertion in *S. cerevisiae.* PCR efficiencies were obtained using qPCR performed three times with four serial 10-fold dilutions in triplicate (Corbett 6000 Rotorgene software).

### Strain construction

dCas9, without fusion to a transcriptional repressor domain, was expressed from a plasmid (# 46920 Addgene (Qi et al., 2013)) under the control of the *TDH3* promoter. The DNA templates for sgRNA production were integrated in place of an exogenous *URA3* cassette immediately downstream of the splice site in the endogenous *snR52* locus using homologous recombination followed by 5-FOA selection. Correct insertion was confirmed by genomic DNA sequencing. *SNR52* is a Pol III-transcribed C/D box small nucleolar RNA (snoRNA) gene and so does not undergo extensive post-transcriptional processing such as capping and polyadenylation, and transcripts from this locus are retained in the nucleus. Additionally, the locus contains a self-splicing site that produces a mature transcript without additional machinery, allowing precise production of a mature sgRNA without any unwanted extensions. Primer sequences for strain construction are listed in Supplementary File 2.

### Yeast growth

The strains used in this study are listed in Supplementary File 3. Strains were grown to mid-log at 30°C in complete synthetic media lacking leucine (for dCas9 plasmid selection). For experiments studying the *GAL1* locus, yeast were grown to mid-log in rich media (YP 2% D/YP 2% Gal) so that the *GAL1* induction kinetics were similar to what we had observed previously (Murray et al., 2012, 2015). dCas9 expression was unaffected by the temporary absence of plasmid selection (Fig. 1 - Figure Supplement 1).

### Yeast growth controls

Assessment of growth in liquid culture (CSM-leucine) was achieved using a Bioscreen spectrophotometer that automatically measures the optical density of cultures at 30°C every 20 min for 24 h. Doubling times were extracted from the gradient of the curves during logarithmic growth.

### Northern blotting

Northern blotting was performed as before (Murray et al., 2012; Nguyen et al., 2014) using asymmetric PCR or *in vitro* transcription with T7 RNA polymerase to generate the radioactive strand-specific probes for *GAL1* and *HMS2* respectively. Primers for these reactions are listed in Supplementary File 4. Northern blots were quantified using ImageJ and images acquired using a Phosphorimager. Images in the figures are scans of exposures to X-ray film.

### Western blotting

dCas9 expression was confirmed after each experiment using an anti-Cas9 antibody (Diagenode C15310258) at 1:5000 dilution and anti-Tubulin (Abcam ab6160) at 1:3333.

### 3’ RACE mapping

Mapping of the 3’ end of the *GAL1* sense transcript was performed as (Nguyen et al., 2014). Primers are listed in Supplementary File 5.

---

**Figure 1 - Figure Supplement 1.**
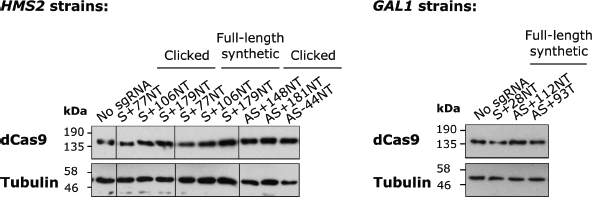
Levels of dCas9 protein in the experimental strains. A Western blot showing levels of dCas9 produced in the strains expressing the indicated sgRNAs against *HMS2* (left panel) and *GAL1* (right panel). Tubulin levels are used as a loading control. Images are taken from the same gels with the same exposure times but with some lanes spliced out (indicated with vertical lines).

**Figure 2 - Figure Supplement 1.**
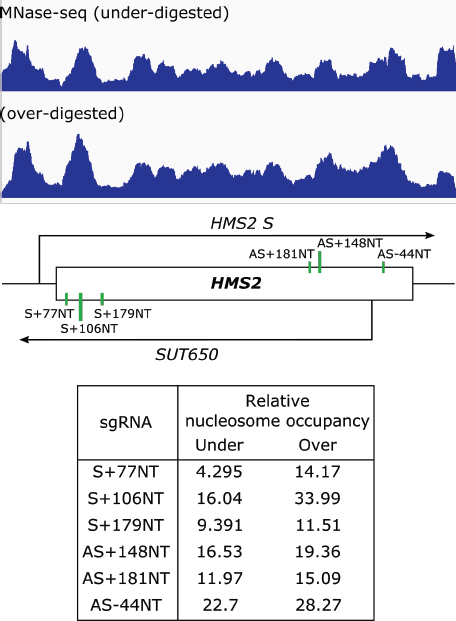
Nucleosome occupancy over the sgRNA target sites does not anti-correlate with level of repression. Nucleosome occupancy map for *HMS2* as measured by sequencing of a sample that has been under- and over-digested with MNase (Knight et al., 2014). The schematic underneath shows the positions of the sgRNA targeting sites. Relative nucleosome occupancy at each sgRNA site is shown in the table for the under– and over-digested samples.

**Figure 3 - Figure Supplement 1.**
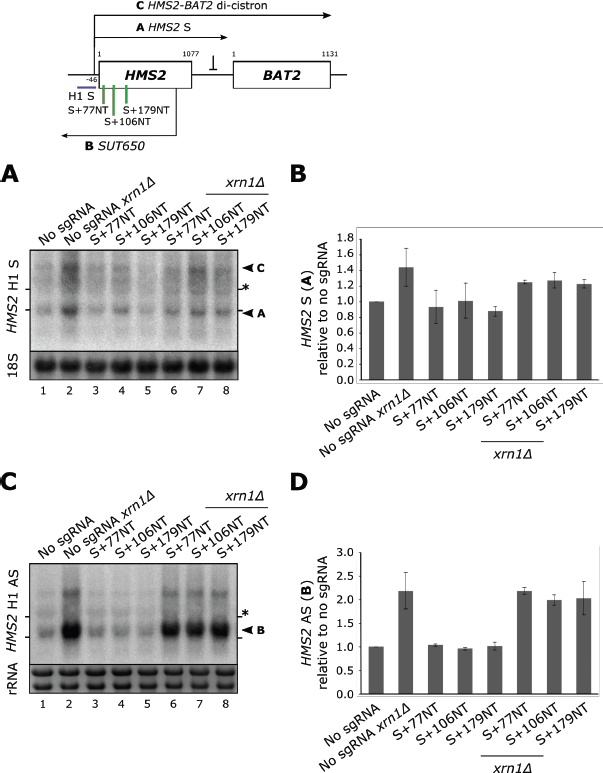
*HMS2* sense-targeting sgRNAs/dCas9 do not affect *HMS2* sense or antisense transcript levels. **A.** A Northern blot probed with *HMS2* sense probe H1 (purple) showing the level of *HMS2* sense (black arrowhead, transcript A) in the no sgRNA control *(snR52::URA3* with dCas9) strain and strains expressing the indicated sense-targeting sgRNAs. Positions of the rRNA are indicated by the short horizontal lines. The 18S rRNA is used as loading control. *represents cross-hybridisation with the 26S rRNA. **B.** Quantification of the Northern blot in (A). N=2, errors represent the SEM. **C.** A Northern blot as (A) but probed with *HMS2* antisense-specific probe H1. Ethidium bromide-stained rRNA is used as loading control. **D.** Quantification of the Northern blot in (C). N=2, errors represent the SEM.

**Supplementary File 1.**
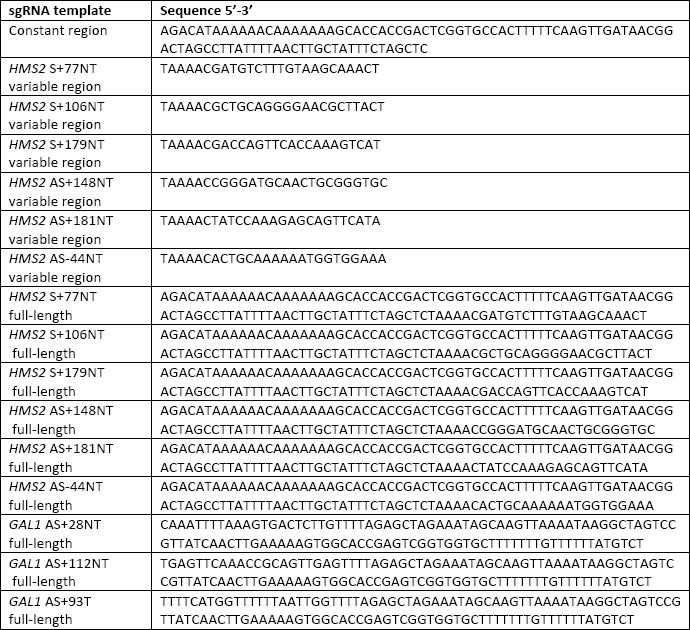
Templates for sgRNAs.

**Supplementary File 2.**
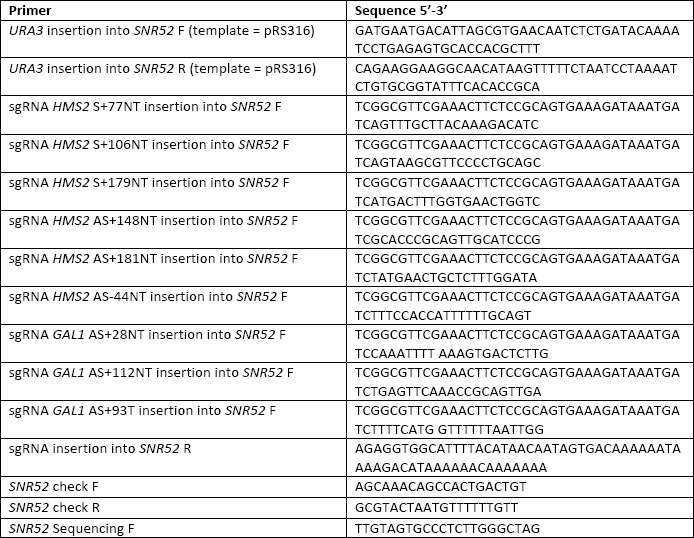
Primers for strain construction.

**Supplementary File 3.**
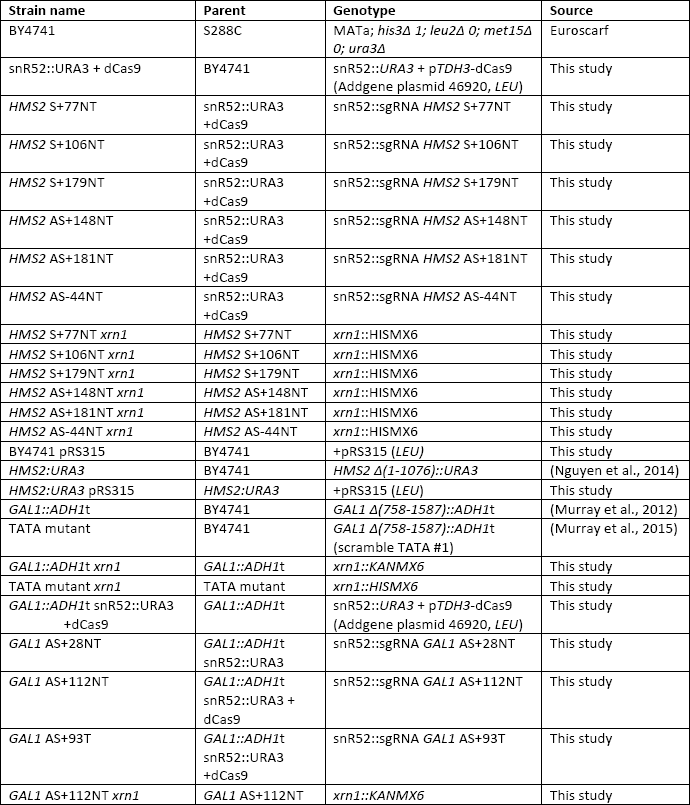
*Saccharomyces cerevisiae* strains used in this study.

**Supplementary File 4.**
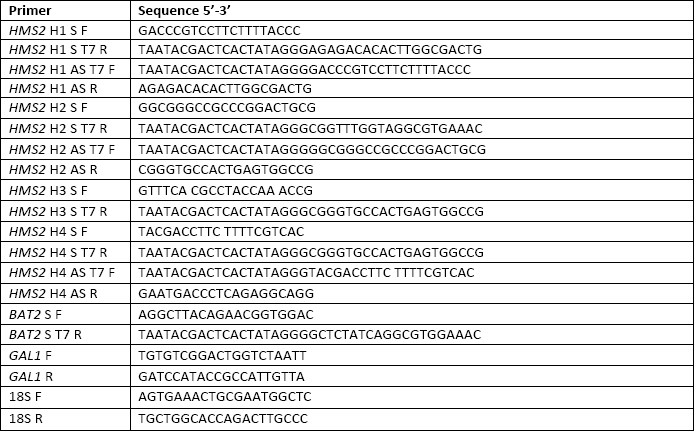
Primers for construction of Northern blot probe templates.

**Supplementary File 5.**
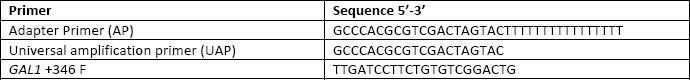
Primers for 3’ RACE mapping of the *GAL1* sense transcript.

